# Implicit motor learning within three trials

**DOI:** 10.1101/2020.04.07.030189

**Authors:** Jennifer E. Ruttle, Bernard Marius ’t Hart, Denise Y. P. Henriques

## Abstract

In motor learning, the slow development of implicit learning is traditionally taken for granted. While much is known about training performance during adaptation to a perturbation in reaches, saccades and locomotion, little is known about the time course of the underlying implicit processes during normal motor adaptation. Implicit learning is characterized by both changes in internal models and state estimates of limb position. Here, we measure both as reach aftereffects and shifts in hand localization in our participants, after every training trial. The observed implicit changes were near asymptote after only one to three perturbed training trials and were not predicted by a two-rate model’s slow process that is supposed to capture implicit learning. Hence, we show that implicit learning is much faster than conventionally believed, which has implications for rehabilitation and skills training.

## Introduction

An established convention of motor learning asserts that automatic or implicit components of learning emerge later in training following an initial more explicit or declarative stage, even for skill-maintenance tasks, like adaptation ^1–6^. Perturbations in reach, saccade and locomotion adaptation evoke relatively quick adjustments to behaviour ^4,6–12^, and some work has attempted to either infer implicit learning based on the assumption that implicit and explicit adaptation simply add to produce behavior ^13^ or measure it in paradigms that require explicitly suppressing natural responses to visual feedback ^11,12^. However, it has not been directly measured how quickly implicit changes emerge. Two main implicit changes involved in error-based learning are updates in internal models as well as the resulting changes in our state estimates ^14–16^. Here, we show that implicit changes during visuomotor adaptation occur immediately and do not require prolonged training at all.

One hallmark of implicit learning: reach aftereffects, is the persistence of motor changes even when the perturbation is removed, which is thought to reflect a change in internal models during adaptation ^15,17^. Yet reach aftereffects are almost always measured after hundreds of training trials, so its time course is largely unknown.

Another implicit change is a shift in our perceived hand location or state estimate that occurs in both visuomotor and force field adaptation ^14,18–22^. A further shift in estimates of hand position can be attributed to efferent-based updates of the internal model ^21,23^. These two sources of hand location estimates have been shown to be unaffected by cognitive strategy and are largely independent ^24,25^. Work from our lab has shown that the changes in hand proprioception following passive exposure to a visual discrepancy, without self-generated movements, can partly account for the resulting changes in reach aftereffects as well ^14,26,27^. Thus, while afferent- and efferent-based estimates of hand position are small, they are robust and contribute to movement planning and reach aftereffects.

It is thought that implicit learning arises slowly with exposure to a perturbation along with explicit components of learning ^1,2,6,13^. Our lab has shown that reach aftereffects and proprioceptive recalibration emerge within 6 trials ^28,29^. In the current study, we push this further by having participants alternate between training and testing trials, while adapting to a 30° rotation, its reversal and then error-clamped trials. Each group of participants performed one type of testing trial that could assess either changes in state estimates or internal models. By probing implicit changes after each training trial, we increase the resolution of measuring implicit changes greatly, and can quantify their rate of change throughout learning.

## Results

### Reach aftereffects

To test how quickly reach aftereffects emerge, 47 participants adapted to an imposed perturbation interleaved with no-cursor test trials. We began by investigating whether these test trials affected reach-training using a control group (N=32) that paused instead. Figure 1A shows the reach training performance for both groups. No-cursor reach trials, and two-rate model fits are shown in figure 1B, and rates of change are listed in table 1.

**Figure 1.**
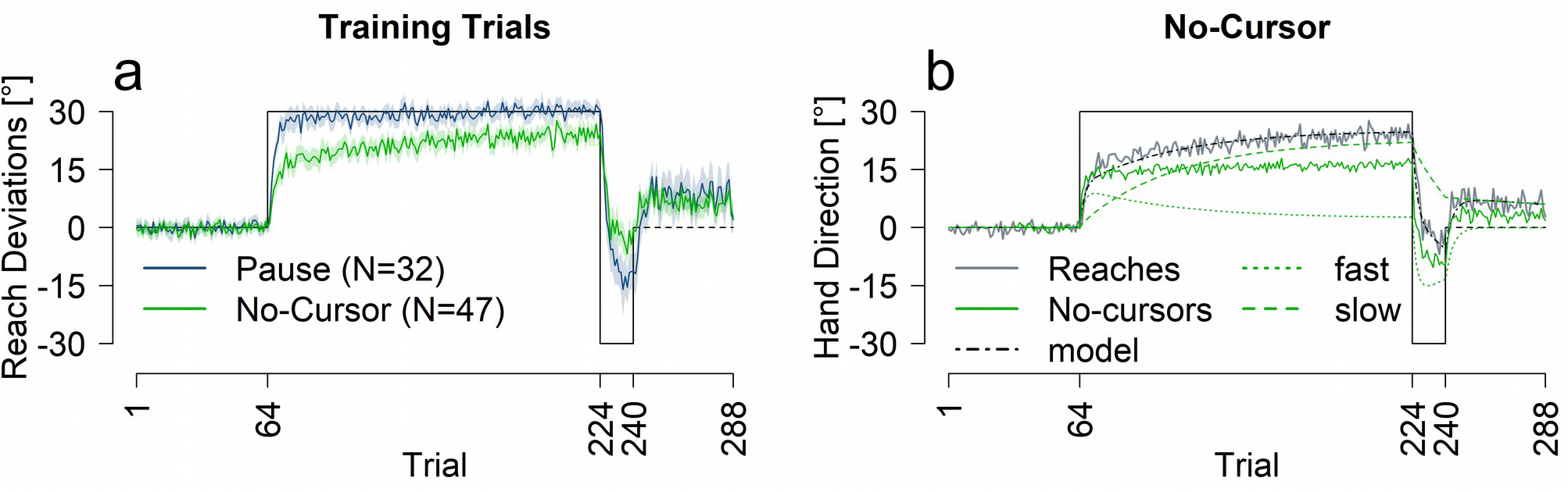
Performance across measures for pause and no-cursor groups. **A:** Reach training performance averaged across all participants for the two groups. **B:** Two-rate model fit (black and green dashed lines), and no-cursor test trials in solid green. For reference, reach performance for the no-cursor group is plotted again in grey. All solid lines are an average of all participants in that group, shaded regions are 95% confidence intervals. Trials included in analysis are as follows: R1=trials 65-68; R1_Late=trials 221-224; R2=trials 237-240; EC=273–288.

**Table 1.** Rate of change and asymptote for reach training trials and measures of implicit learning, for each of the experimental groups, calculated for the first rotation. Reach training trials are represented in the first three rows, then the slow process predicted by the model, with the final three rows being the implicit measure test trial associated with that training group. Averages with 95% CI are reported for all values.

| Rate of change |  |  |  |  |  |
| --- | --- | --- | --- | --- | --- |
|  |  | No-cursor | Pause | Active | Passive |
| Reach training | rate of change | 13.8%<br>[10.7% - 17.7%] | 36.5%<br>[28.3% - 49.8%] | 15.4%<br>[10.8% - 21.8%] | 27.0%<br>[21.4% - 34.8%] |
|  | asymptote | 23.2°<br>[22.0° - 24.3°] | 29.1°<br>[28.5° - 29.7°] | 24.7°<br>[22.7° - 26.2°] | 28.6°<br>[27.8° - 29.5°] |
|  | saturation trial | 21<br>[16 - 27] | 9<br>[6 - 12] | 16<br>[11 - 23] | 12<br>[9 - 15] |
| Slow process | rate of change | 3.4%<br>[3.1% - 3.8%] | 2.8%<br>[2.5% - 3.2%] | 2.1%<br>[1.9% - 2.4%] | 3.5%<br>[3.0% - 4.1%] |
|  | asymptote | 21.4°<br>[19.8° - 23.0°] | 27.2°<br>[25.5° - 28.6°] | 26.0°<br>[23.9° - 27.8°] | 25.2°<br>[22.6° - 27.4°] |
|  | saturation trial | 74<br>[67 - 82] | 96<br>[85 - 109] | 117<br>[107 - 130] | 64<br>[54 - 74] |
| No-cursor or localization | rate of change | 56.9%<br>[27.4% - 58.5%] | - | 93.9%<br>[51.3% - 95.7%] | 100%<br>[29.0% - 100%] |
|  | asymptote | 15.3°<br>[13.8° - 16.9°] | - | 9.8°<br>[8.1° - 11.6°] | 6.9°<br>[5.7° - 8.2°] |
|  | saturation trial | 3<br>[1 - 3] | - | 1<br>[1 - 3] | 1<br>[1 - 6] |

The time course of adaptation has been described by a state-space model that includes two processes, a process with slow learning and forgetting and a process with fast learning and forgetting ^30^. The model’s fast and slow processes have been suggested to map onto explicit and implicit components of learning respectively ^13^. While not the main focus of this study, here we compare the rate of change of the models’ slow process with that of actual measures of implicit learning.

We find changes in reach aftereffects and state estimates to occur much faster than expected. State estimates asymptote after a single trial and are well described as a proportion of the perturbation. Our results challenge the convention that implicit learning is slow, and show that some implicit changes emerge before, and likely independently and not inferable from, explicit changes in motor control.

#### Training Trials

To investigate whether the type of intervening test trial affects training performance (Fig 1A), we conducted a mixed ANOVA with group (no-cursor or pause) and trial set (R1, R1_Late, R2 and EC, described in figure 1 & 6). We found an effect of trial set [F(3,186)=415.30, p<.001, η2=.82] and an interaction between trial set and group [F(3,186)=11.78, p<.001, η2=.11]. The interaction seems to be driven by the slower learning and much smaller rebound in the no-cursor paradigm. Follow-up t-tests show a significant difference between the pause and no-cursor group during R1, R1_late, R2 and EC trials sets with p<.01. We fit the two-rate model to the averaged reach deviations for each group (see figure 1B and table 2). The model predicts average performance well for the pause group and reasonably well for the no-cursor group. The smaller learning rate parameter values for the no-cursor group versus the pause group (table 2), are mimicked in the rates of change (table 1), which may be explained by slight interference from the no-cursor trials. Nevertheless, the model fits warrant comparison between reach aftereffects and the model’s slow process.

**Table 2.** Model parameters and goodness-of-fit estimates. All twoRate AIC’s are smaller than respective oneRate AICs indicating a better model fit from a two-rate model. Relative likelihoods below .05 are bolded. Parameter values could vary between 0 and 1, inclusive.

| Group | Rs | Ls | Rf | Lf | MSE | Two-Rate AIC | One-Rate AIC | One-Rate likelihood |
| --- | --- | --- | --- | --- | --- | --- | --- | --- |
| No-Cursor | 0.991 | 0.036 | 0.737 | 0.148 | 3.061 | 13.901 | 19.11 | 0.074 |
| Pause | 1 | 0.055 | 0.825 | 0.226 | 8.345 | 19.293 | 25.697 | **0.041** |
| Active Localization | 0.999 | 0.03 | 0.76 | 0.158 | 4.053 | 15.605 | 27.004 | **0.003** |
| Passive Localization | 1 | 0.054 | 0.74 | 0.236 | 7.425 | 19.548 | 27.207 | **0.022** |

#### Testing Trials

We found reach aftereffects were present at the first trial set, after only 1-4 rotated training trials, compared to those of no-cursors from the aligned phase [t(46)=20.97, p<.001, d=4.12, η2=.82, 10.94°]. Using an exponential decay model (see methods) we calculated a rate of change and the trial at which the no-cursor deviations are at asymptote, focusing on the first rotation. We found reach aftereffects had a rate of change of 56.9% (CI: 27.4% - 58.5%), attaining asymptote within 3 trials (see table 1 for all rate of change values). Both the substantial rate of change and, early trial at which changes saturate in no-cursor reaches, i.e., the implicit reach aftereffects, show that implicit adaptation develops rapidly. In fact, aftereffects saturate well before reach training does (56.9% > 13.8%, 95% CI: 10.7% - 17.7%), which asymptotes only at the 21st trial for this group and 9th trial for the control, pause group.

Furthermore, the rate of change of the slow process (3.4%) is much lower than the rate of change of the reach aftereffects (56.9% > 3.4%, 95% CI: 3.1% - 3.8%). Additionally, fitting the two-rate model’s slow process to the reach aftereffects, to see if the model’s output still matches reaches increases the AIC from 13.9 to 29.17 significantly decreasing the fit (relative log-likelihood: 0.0005; see OSF https://osf.io/9db8v/ for details). This all shows the rate of change in reach aftereffects is much higher than what would be expected of a slow, implicit process, or indeed that of the two-rate model’s slow process.

### Hand localization shifts

An additional 64 participants adapted to the same perturbation schedule, interleaved with test trials that measured estimates of the hand location after the trained hand was displaced by a robot manipulandum (passive localization, N=32, Fig 2A&D) or by the participant themselves (active localization, N=32, Fig 2A&C). We once again used the pause group as a control (Fig 2A).

**Figure 2.**
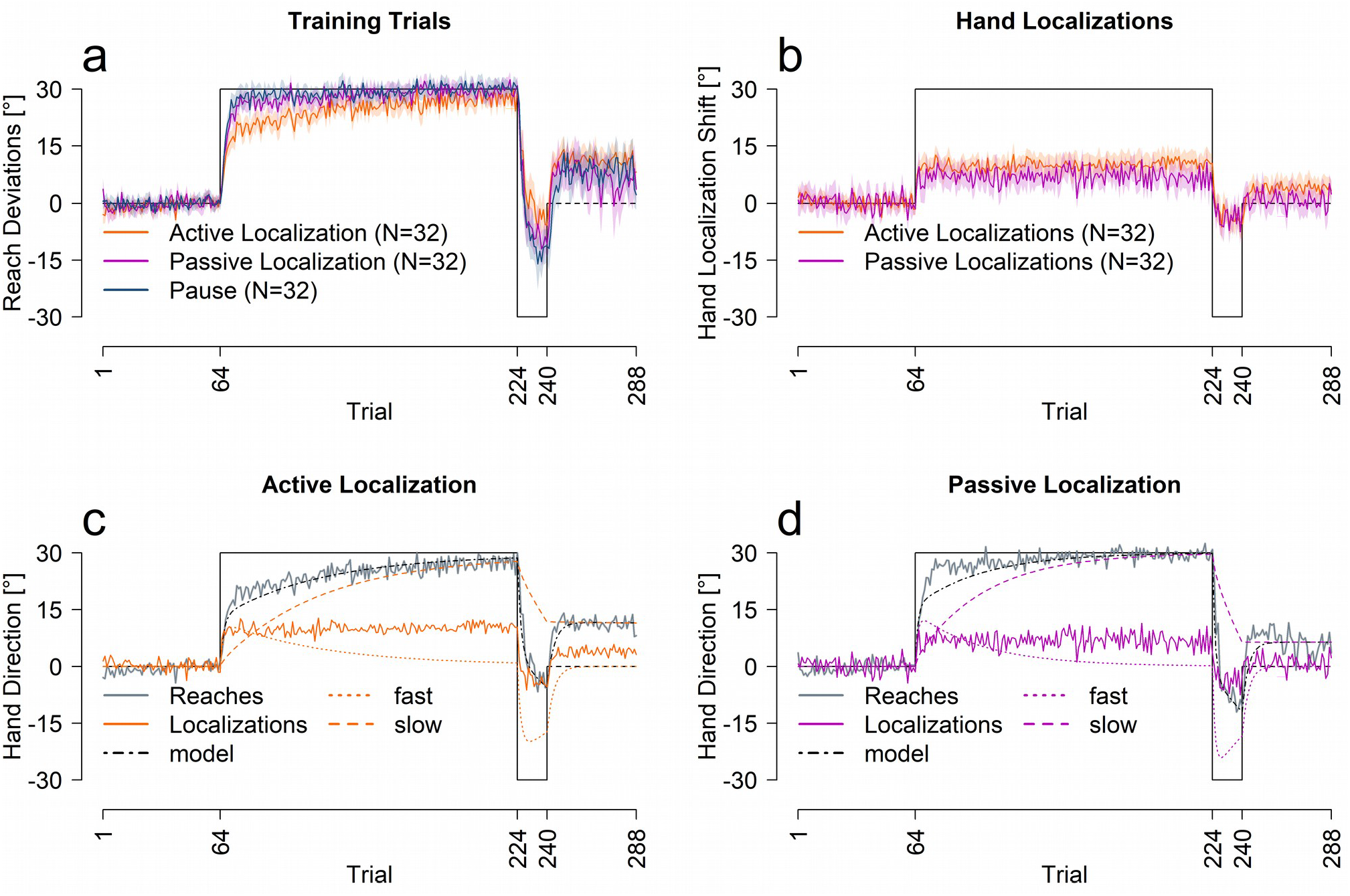
Performance across measures for passive, active and pause groups. **A:** reach training performance averaged across all participants for each corresponding group. **B:** Hand localization performance for the two groups. **C&D:** Model predictions for the active and passive localization groups. All solid lines are an average of all participants in that group, shaded regions are 95% confidence intervals.

#### Training Trials

We conducted an ANOVA with the same factors as the previous experiment, trial set and group (active localization, passive localization and pause) on training performance (Fig 2A). As expected, reach deviations varied across trial set [F(3,279)=537.99, p<.001, η2=.80], and there was a significant interaction between trial set and group [F(6,279)=8.29, p<.001, η2=.11], but no effect of group on its own [F(2,93)=1.90, p=0.15]. Follow-up ANOVAs show that learning was slower in active localization compared to the other conditions [all p<.001]. We also fit the two-rate model to the reach data from the passive and active localization groups, shown in figure 2C&D. The model fits the reach data well, but as shown in table 2, the learning rates for the active group are slightly lower than the passive or pause group. Importantly, the retention parameters are very similar across all three groups, indicating the same ability to retain what was learned. In summary, despite a small effect of test-type, the two-rate model explains the reach data well.

#### Test Trials

We also compare the time course of changes in estimating the location of the unseen, adapted hand across training: for the passive vs. active localization shown in Fig 2B. Estimates of hand position show a rapid shift on the first trial after the initial perturbation is introduced for both active (8.95°) and passive localizations (6.46°). These shifts do not increase with further training with both groups achieving 93.9% and 100% of asymptote within one rotated training trial. Seeing as changes in state estimates of hand location appear incredibly fast, these localization shifts can not follow from motor adaptation. Instead, they directly result from the perturbation in a single trial.

Despite similarly quick shifts in hand localization (Fig 2B), a mixed ANOVA revealed a significant difference in hand estimates between the active and passive localization groups [F(1,62)=6.28, p=0.014, η2=.05], across trial sets [F(3,186)=96.97, p<.001, η2=.43] and an interaction between trial set and group [F(3,186)=2.93, p=0.04, η2=.02]. Follow-up t-tests indicate larger shifts in felt hand position in the active localization group both during the initial [t(51.43)=2.37, p=0.028, d=.59, η2=.08, 2.92°] and final [t(61.78)=2.98, p=.016, d=.74, η2=.11, 4.35°] trial set of the first rotation and at the end of the error clamp phase [t(61.99)=2.73, p=.016, d=.68, η2=.11, 3.5°]. Thus, even though the participants in the active localization group adapted their cursor movements slightly slower than the passive group, the active group showed a slightly larger shift in felt hand position, that didn’t quite reach significance for the counter rotation [t(58.93)=−0.15, p=.88]. This small separation between active and passive localization shifts reflects the updates in the predicted sensory consequences that further shift active hand localization compared to passive ^19,23,27^.

More importantly, Fig 2C&D, show that these changes in state estimates are not captured by the slow process. To test this, we used the same, simple exponential decay model to quantify the rate of change in the passive and active localization data as well as in the models’ slow processes. The rates of change are much higher for the localization data than the respective slow process [Passive: 3.4% < 100%, 95% CI: 29.9% – 100%; Active: 2.1% < 93.9%, 95% CI: 63.2% - 100%]. Thus, shifts in hand localization in no way resemble the observable pattern of the slow process.

Given that the rate by which estimates of hand position changes do not match the slow process and saturate in 1 trial (see table 1), we next tested whether these shifts could simply be described as a proportion of the perturbation. When the changes in hand position were fit with a linear regression estimating the proportion of the perturbation accounted for, we see a reasonable fit between actual hand location estimates and these simple models (see Fig 3A-B). The fitted slopes are consistent with previous studies, where the change in felt hand position was usually 20-30% of the perturbation ^19,31,32^. Figure 3D&E shows a one-parameter model (black line) that estimates the size of the average localization shift as a proportion of the perturbation using all trials (colored lines). Even though the model over-estimates the size of change in hand localizations during the reversal period, it is clear that a step-wise function is a better fit than an exponential, further providing support for the conclusions that hand localizations shift incredibly fast, too fast to be the slow process in the two-rate model. The small over-estimate of this fit for reversal phase in the active localization group reflects what is a relatively slower rate of changes for this phase (for all measures, but not the slow process), which could partly reflect retrograde interference. Nonetheless, the change in localization during the reversal seems to occur even prior to the change in cursor-reach for this reversed shift of perturbation (compare curves within Fig 2D and 2E). Given that hand localization shifts occur before other changes, they are able to guide subsequent motor adaptation processes.

**Figure 3.**
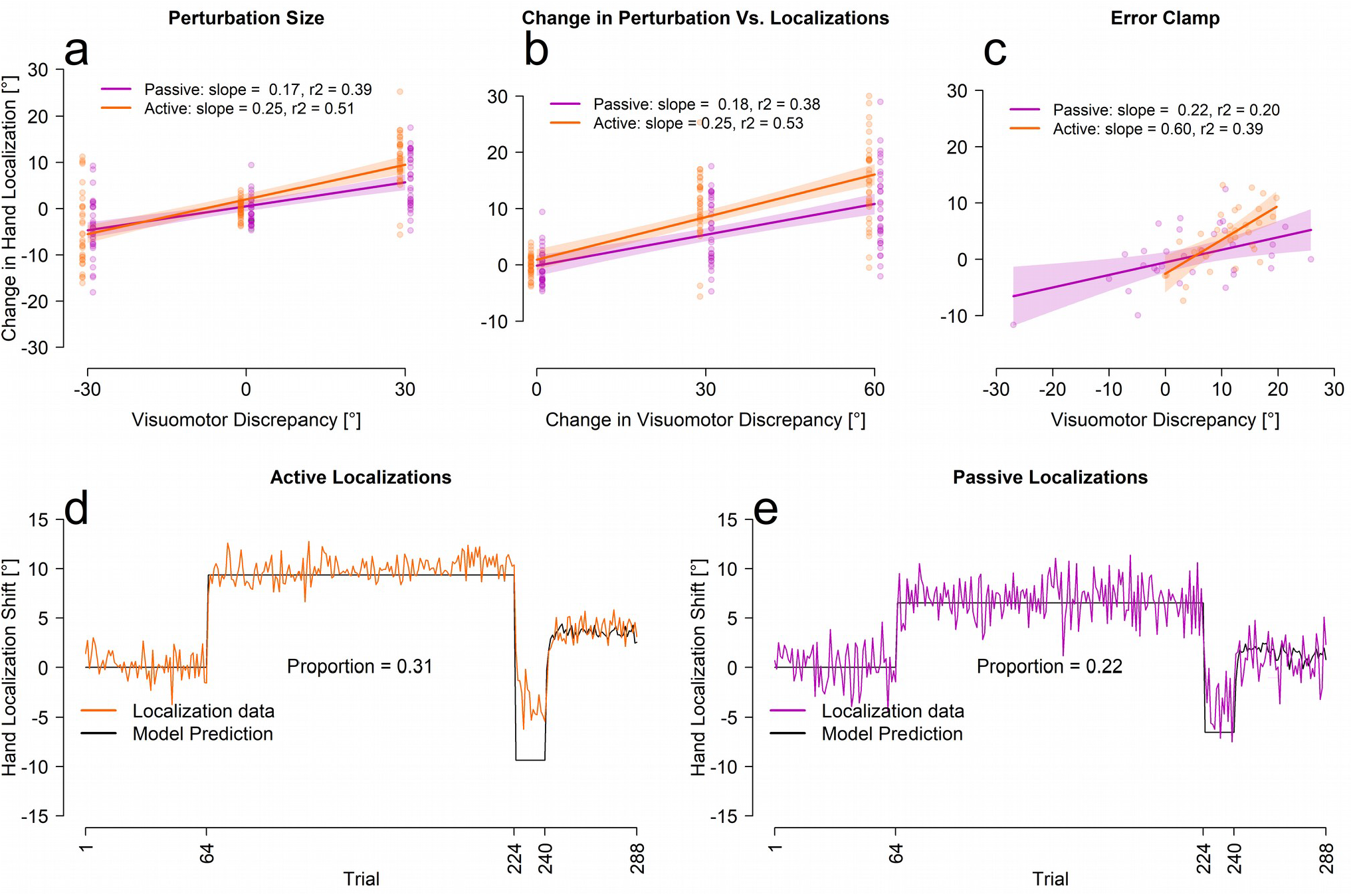
Linear and proportional fits between localization and error clamp trials. **A-C:** Fits between localizations and either the size of the visual discrepancy (aligned = 0, first rotation = 30, second rotation = −30), the absolute change in size of visual discrepancy (aligned = 0, first rotation = 30, second rotation = 60) and the participants average performance on the final 16 error clamp trials. The shaded regions represent the 95% confidence interval around the regression line. The last four initial rotation trials, the last four reversal trials and the last four aligned training trials were used for figures A&B. **D-E:** The proportional models’ prediction and the averaged participant performance for each localization test trial separately.

### Implicit Measures of Learning

As both reach aftereffects and hand localization shifts are implicit and potentially driven by similar processes we compared the first test trial after the first rotated training trial and found no significant difference between the reach aftereffects, and either of the active [t(73.54)=−1.08, p=.28] or passive [t(59)=−1.81, p=.08] localization test trials. This finding is consistent with the relatively small correlations between angle at peak velocity during error clamp trials and the corresponding estimates of hand location seen in figure 3C. We speculate that these initial reach aftereffects may mainly reflect the changes in hand location estimates, or a similar training signal ^14,19,26^, before additional sources of information emerge to create even larger shifts in reach aftereffects but no further shift in hand localizations.

### Speed of Learning

While tangential to the main goal of this study, we found that intervening trials that involve active movements (no cursor, or active localization where participants moved their own trained hand) slowed down learning when compared to just passive hand displacements or a pause in time. We can in fact predict for individual participants whether they made self-generated movements in the interleaved testing trials or not for 118 / 143 participants (82%; chance=79/143 or 55%; p<.001 binomial exact test) based on a multiple logistic regression model without interactions, using the parameters of the two-rate model as predictors. The exponential decay model also predicts a much slower rate of change for reach training when the test trial involves an active movement (no-cursor: 13.8%, active localization 15.4%) compared to when it does not (pause: 26.5%, passive localization 27.0%). This shows that learning is slowed by having active intervening movements made in the absence of visual feedback. This means that our measures of the rate of change may be underestimating the real speed of implicit learning in some of the processes.

## Discussion

While many studies measure and model the time course of reaches in response to a perturbation ^13,33^, very few investigate the emergence of other outcomes of training, such as reach aftereffects and changes in estimates of hand position, but see: ^28,29,34^. In the current study, we measure implicit changes as reach aftereffects and estimates of the passively and actively displaced hand’s position, at high temporal resolution. This is accomplished by following every reach training trial that has aligned, rotated or error-clamped cursor feedback with one test trial. We find that reach aftereffects and changes in estimates of hand position emerge and saturate rapidly, within 1-3 trials of visuomotor adaptation training (see Fig 4). That is much faster than when reach adaptation saturates (9 trials at best), let alone the two-rate model’s slow process (64 trials at best). This suggests that implicit changes do not follow explicit changes and play an important role in initial learning. Indeed, changes in hand-localization are so fast, they can best be explained as a proportion of the visual-proprioceptive discrepancy experienced on the previous trial. In sum, given that our measures of implicit learning saturate within 1-3 trials, implicit learning can hardly be characterized as “slow”.

**Figure 4.**
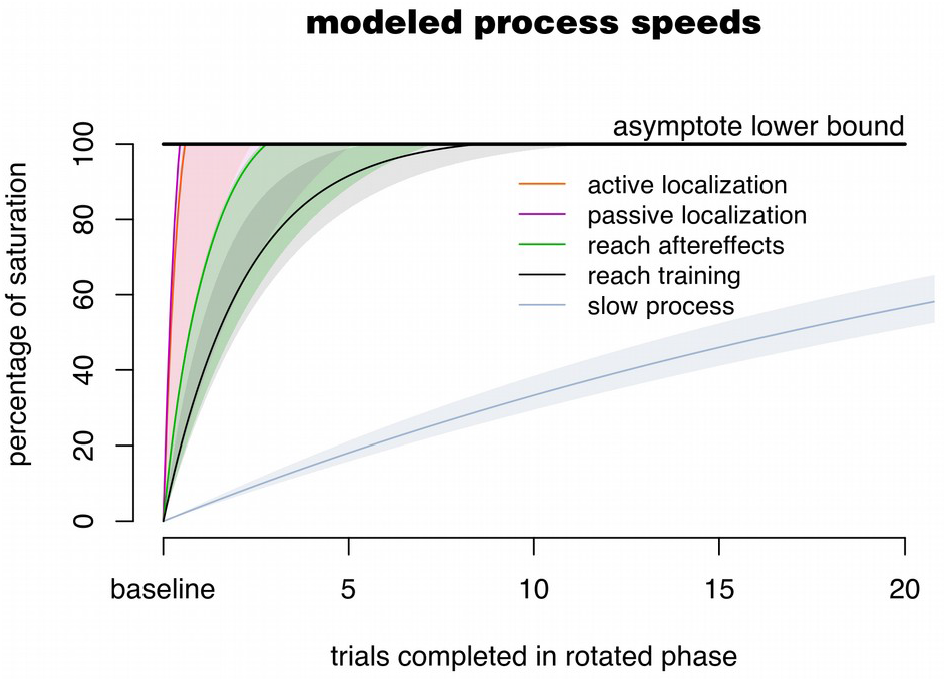
Overview of Results. For each of the five processes considered in the paper, the plot shows the change from baseline (0%) to the lower limit of the 95% confidence interval of the asymptote (100%). For the reach adaptation and the two-rate model’s slow process, the highest rate of change is used. As is clear, the measures of implicit adaptation (reach aftereffects and hand localization shifts) are faster than adaptation and much faster than the model’s slow process.

As expected, reach performance with a rotated cursor for all four groups adhered well to a model that consists of a fast and a slow process ^13^. Adaptation, and hence implicit processes in other studies are sometimes slower than what we found for our four groups, especially in the case where targets span the entire radial workspace ^13^. However, given the fact that learning-induced changes in hand estimates and reach aftereffects saturate within one to three trials, i.e. before experiencing all four targets, it’s unlikely that this saturation would have been much delayed or reduced if we tested more locations. Regardless, all other processes would likely have slowed down as well, so that our direct measures of implicit adaptation would still be faster. Of note is also that while we found that some of the interleaved testing trials slowed down adaptation, the underlying implicit processes were still very fast.

Estimates of hand location are incredibly quick to shift, with participants only having to experience one rotated training trial to elicit the full shift. Hand localization responses are thought to measure the brain’s state estimate of hand position and likely rely on at least two signals: an efferent-based predictive component and an afferent-based proprioceptive component, that both change during visuomotor rotation training ^23,35^. Active localization reflects both, and indeed exhibits slightly larger shifts than passive hand localization, which is consistent with previous findings ^23,27^, as is the size of the shift in hand localization of 20-30% of the rotation ^18,29,31^. We see here that a proportional fit seems to explain changes in hand estimates throughout the adaptation task, especially during the error-clamp phase where the size of the visual-proprioceptive discrepancy is determined by the size of the reach deviation. This is essentially a step-function which indicates that the process of changing estimates of hand location is qualitatively different from other processes of motor learning.

Reach aftereffects also emerge incredibly quickly, while not reaching asymptote as fast as shifts in hand localization. The similar size of aftereffects and hand localization shifts after just one rotated training trial potentially indicates a shared source. In addition, participants who perform no-cursor reaches with minimal instruction or more detailed instruction (to ensure strategy wasn’t used) show similar rates and extents of learning of reach aftereffects (see OSF; https://osf.io/9db8v/), which is in line with some previous findings ^24,36^. If no-cursor reach deviations reflect implicit changes in state estimation, these arise much quicker than previously thought bolstering recent claims that the earliest wave of muscle activity during adaptation is influenced by implicit motor learning ^37^.

Other work on the time-course of implicit adaptation uses primarily two approaches. Either implicit adaptation is indirectly inferred from reach deviations and a measure of explicit learning ^3,13^, or it uses error-clamped feedback paradigms ^12^. Results from such approaches indicate that implicit learning is slow, and we can only speculate here about why our findings are so different. In the first approach, subtracting a measure of explicit adaptation from training reach deviations relies on a largely untested assumption that implicit and explicit adaptation linearly add to produce behavior ^13,30^. Aside from the effect the aiming task may have on adaptation ^38,39^, this is not a biologically plausible mechanism, and it should not be surprising that an actual measure of implicit adaptation, as we use here, shows a different time course. Our results set the nature of the mechanism by which implicit and explicit adaptation are combined as a topic for future study.

In the second approach for measuring implicit adaptation, which uses error-clamped feedback to get at the time course of implicit adaptation, participants are instructed about the nature of the paradigm, prompted to ignore the visual feedback, and faced with unnaturally smooth reach trajectories throughout the task ^12,40^. Since this context necessarily increases external error attribution it should also suppress implicit learning ^41^. This could explain why error-clamped feedback paradigms slow down implicit adaptation compared to how it naturally occurs here. In sum, both approaches to assessing implicit adaptation have drawbacks, that straightforward interleaving of trials doesn’t have, although our method does show some interference, potentially slowing down adaptation processes. It will remain an interesting challenge to unify results from all paradigms.

The results here raise a few other questions. We observe that reach aftereffects are as large as hand localization shifts after 1 trial of perturbed feedback, which may be an indication that hand localization shifts contribute to reach aftereffects. More compelling evidence comes from previous studies where a strong correlation between these measures was found ^24,26^. We have shown that the size of both proprioceptive and predictive components of hand localization shifts can predict separate components of reach aftereffects ^41^. However, all evidence that hand localization shifts contribute to reach aftereffects is correlational, so that the mechanism remains unknown. In addition, while here we did manage to look at the time-course of implicit processes, the relevance for rehabilitation and skills training would lie in how these consolidate, as eventually we need to be able to move without exerting explicit control. That as little as 5 trials suffice for savings ^42^, is hopeful and in line with the high speed of implicit processes we find here.

## Conclusion

We show here that the conventionally implicit components of motor learning; no-cursor reach deviations, and changes in estimates of hand location emerge very rapidly. The fast emergence of reach aftereffects and changes in hand estimates indicate implicit components of motor learning appear before or alongside explicit components of learning. Perhaps some implicit processes lead or maybe drive motor learning, unlike previously believed, as certainly they do not lag behind explicit processes. In addition, our results provide further evidence that implicit learning consists of at least two sub-processes that separately contribute to adaptation, and that both can be extremely fast.

## Methods

### Participants

143 (mean age=20.31, range=17-46, females=101) right-handed, healthy adults participated in this study. All participants gave written informed consent prior to participating. All procedures were in accordance with institutional and international guidelines. All procedures were approved by the York Human Participants Review Subcommittee.

### Apparatus

The experimental set-up is illustrated in Figure 5. While seated, participants held a vertical handle on a two-joint robot manipulandum (Interactive Motion Technologies Inc., Cambridge, MA, USA) with their right hand such that their thumb rested on top of the handle. A reflective screen was mounted horizontally, 14 cm above the robotic arm. A monitor (Samsung 510 N, 60 Hz) 28 cm above the robotic arm presented visual stimuli via the reflective screen to appear in the same horizontal plane as the robotic arm. A Keytec touchscreen 2 cm above the robotic arm recorded reach endpoints of the left hand, to unseen, right hand targets (see ^18^ for more details). Subject’s view of their training (right) arm was blocked by the reflective surface and a black cloth, draped over their right shoulder. The untrained, left hand was illuminated, so that any errors in reaching to the unseen, right target hand could not be attributed to errors in localizing the left, reaching hand.

**Figure 5.**
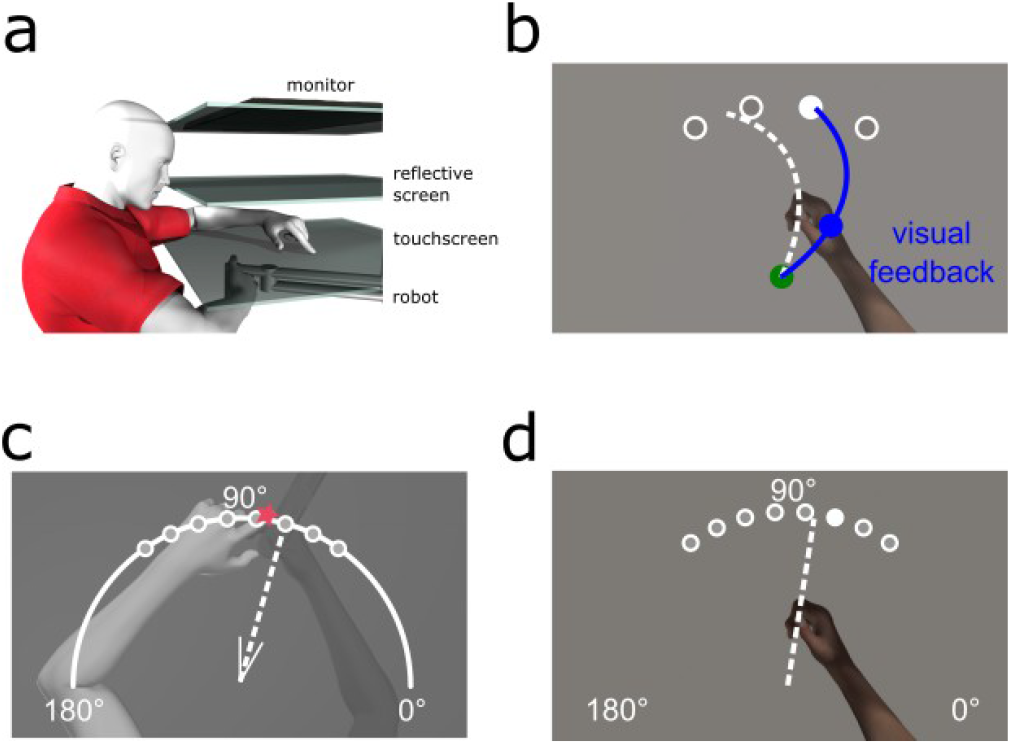
Experimental setup and design. **A:** Side view of the experimental set-up. The top layer is the monitor, middle layer is the reflective screen, and the bottom opaque layer is the touchscreen. The robot is depicted beneath with the participants’ right hand grasping it. **B-D:** Top views of task specific set-ups. **B:** Training (and Clamp) trial. The home position is represented by a green circle with a 1 cm diameter; located approximately 20 cm in front of the subject. Targets are represented by white circles with a 1 cm diameter located 12 cm radially from the home position at 60°, 80°, 100° and 120°. Participants hand cursor was also a 1 cm diameter blue circle. **C:** Localization test trial. Participants were either passively moved to one of the eight target locations, or actively moved their hand in the direction suggested by the white wedge, consisting of two short straight lines (V-shaped) at the home position, these real and suggested locations are 55°, 65°, 75°, 85°, 95°, 105°, 115° and 125°. Subsequently, participants used a touch screen to indicate on a white arc spanning 180° where their unseen right hand was. **D:** No-cursor test trial. Participants made ballistic reaches to one of the 8 target locations also used in localization without any visual feedback of their movement. Figures were made using Poser Rendering Software version 11, https://www.posersoftware.com/.

### Trial Types

#### Reach-training trials

Participants, regardless of group, reached as accurately as possible with their right hand to one of four possible target locations, 60°, 80°, 100° and 120°, which were shown once in a cycle of four trials before being repeated (see figure 5B). In all reaching trials, i.e., with cursor, with clamped cursor and with no cursor, participants had to reach out 12 cm from the home position to a force cushion within 800 ms. Participants received auditory feedback throughout training indicating if they met the distance-time criteria or not. The target would then disappear, and the robot manipulandum returned the right hand to the home position where they waited 250 ms for the next trial. The hand cursor was aligned with the hand for the first 64 training trials, then rotated 30° CW for 160 training trials and then rotated 30° CCW for 16 training trials. This was followed by 48 error-clamped trials, dashed lines in Fig 6, which were identical to the reach training trials except that the cursor always moved on a straight line to the target. The distance of the error-clamped cursor from the home position was identical to the distance of the hand from the home position.

**Figure 6.**
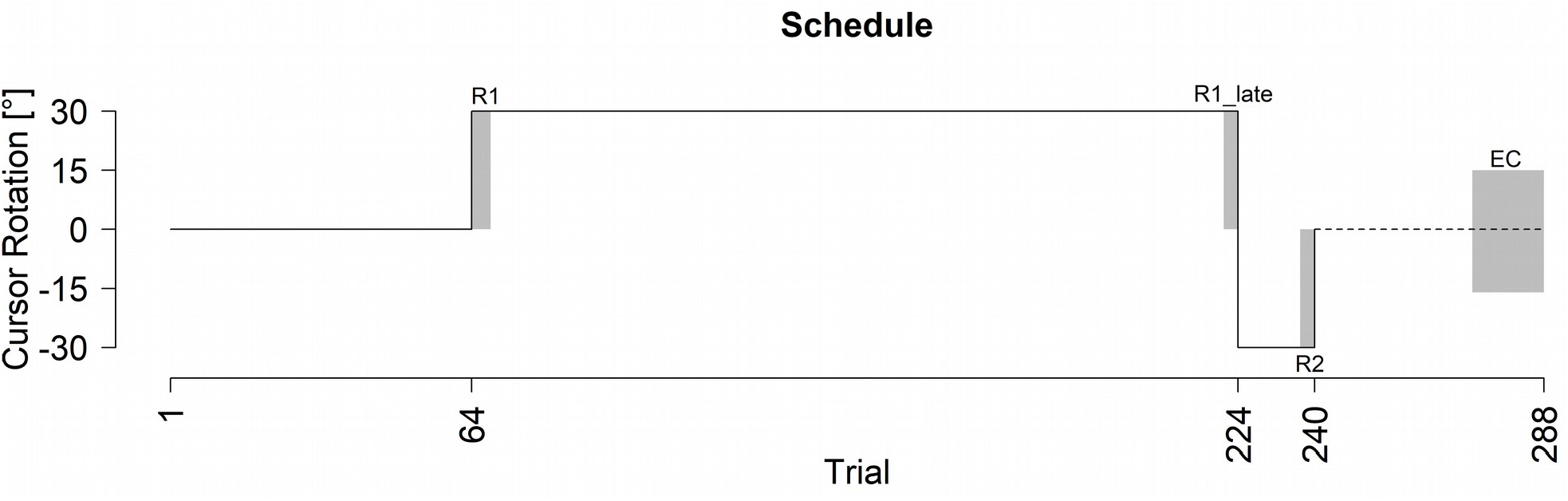
Experimental Schedule. Participants reached to visual targets with a perturbation denoted by the black line. The dotted line at the end of the paradigm signifies clamp trials where there was no visual error as the cursor always moved to the target regardless of the participants movement direction. Trials included in analysis are as follows: R1=trials 65-68; R1_Late=trials 221-224; R2=trials 237-240; EC=273–288.

#### Test trials

Participants alternated between one training trial, as described above, and one of four possible test trials. That is, test trials are interleaved with training. Each type of test trial was performed exclusively by participants in one group. These test trials were: (1) a no-cursor reach to a target, “No-cursor”, N=47, (2) a short pause phase with no hand movement, “Pause”, N=32, serving as a control group, (3) localization of the unseen hand position when the hand was passively moved by the robot, “Passive localization”, N=32, or (4) localization of the unseen hand after it was actively moved by the participant, “Active localization”, N=32. After each test trial, the robot returned the participants’ hand back to the home position. Test trials were always executed to one of two targets, 5° on either side of the training target, to reduce distance between test and training targets. All eight targets (55°, 65°, 75°, 85°, 95°, 105°, 115° and 125°; one on each side of each of the training targets) were cycled through before being repeated.

#### Reaching without a cursor

For the no-cursor group, their test trial required reaching out, again 12 cm, to one of the eight test targets (Fig 5D) without a cursor representing their hand. The same distance-time criteria as in reach-training applied but without reinforcing sounds. This group originally had 32 participants who were simply told that there would be no cursor for these trials. We later add 15 more participants who were specifically told not to include any learned strategy, similar to a previous study in our lab that used a PDP technique and showed no explicit component for a 30° rotation 24,25. Since the results did not differ between these two sub-groups, (see OSF for details: https://osf.io/9db8v/), the results were collapsed for analyses. Our results are consistent with the current idea that implicit learning caps around 15° ^12^ and previous studies which found no difference in the size of reach aftereffects when participants are not told about the rotation, then during a set of no-cursor trials, either include or exclude any strategy developed in the training trials ^24,25,36^.

#### Hand localization

The two hand localization groups did test trials measuring estimates of unseen hand location in order to assess different components of state-estimation, after every training trial. For both localization trials (Fig 5C), a white arc would appear on the screen, spanning from 0° to 180°, the arc was 12 cm away from the home position. Then the hand was either passively displaced by the robot to one of the eight target locations (passive localization) or the hand movement was self-generated by the participant (active localization). Passive movement of the hand took 650 ms to cover the 12 cm distance. In active localization trials, participants chose their own hand-target location. They were guided with a small V-shaped, 30° wedge that appeared at the home position, the middle of the V-shaped wedge was oriented to the passive localization targets. This active, self-generated movement was stopped by a force cushion at the 12 cm mark. Regardless of localization trial type once their right, unseen target hand was locked in place, participants used their visible, left index finger, to indicate on the touchscreen, along a 180° arc, where they believed their right, stationary, unseen hand was. The arc was continuously visible until the touchscreen registered the participants estimate.

### Data Analysis

We analyzed the reach-training for the no-cursor group and the two hand localization groups separately using the pause group as a control. The reach training trials, hand localization trials and no-cursor trials were analyzed separately from each other, but their rates of change (see below) can be compared.

#### Reaching with a cursor and clamp trials

To quantify reach performance during training, the angular difference between a straight line from the home position to the target and a straight line from the home position and the point of maximum velocity is computed.

#### Hand Localization

Estimates of hand location in both the passive and active localization groups were based on the angular endpoint error between the movement endpoint of the right unseen hand and the left hands responses on the touchscreen.

#### Reaching without a cursor

To determine if participants exhibit reach aftereffects as a result of training, we measured reach endpoint errors during no-cursor trials. The reach error is calculated based on the angular deviation between the reach endpoint and the target location, relative to the home position. We used the endpoint error, instead of maximum velocity to be able to compare no-cursor trials to hand localization trials. However, a comparison between no-cursor reach deviations at endpoint and at maximum velocity is included in the R notebook.

### Analyses

All data was visually screened for incorrect trials. Subsequently, outliers of more than three standard deviations across participants within each trial were also deleted. In all, 2.2% of the data was removed. One participant had to be removed from the no-cursor instructed group as they did not complete the task appropriately. All measures were normalized, by subtracting out each subjects’ average performance during the second half of the aligned session (e.g. trials 33-64). To see if there were changes in training and test trials, we conducted ANOVAs consisting of a within-subjects factor of trial set and a between-subjects factor of group. The trial-set factor consisted of four levels: the first 4 rotated trials (R1), the final 4 trials from the first rotation (R1_Late), the final 4 trials from the second rotation (R2) and the last 16 trials, to allow for a less noisy estimate, from the clamp phase (EC). All analyses ignored target location, but each bin of four trials contains a trial to each of the four training targets (effects at different target angles are not distinguishable, see the R notebook). Significant main effects and interactions were followed-up by pairwise comparisons. All results are reported with a Welch t-test and an alpha of .05, where necessary with an fdr correction applied using the p.adjust function in R.

### Two-Rate Model

We fitted the two-rate model ^30^ to our data. This two-rate model is composed of a slow process that slowly increases over time until it is the driving force of performance, and a fast process that rises quickly but eventually decays back to zero. The sum of these two processes determines the overt behaviour and can explain the rebound seen in the error-clamp phase. During error-clamps, neither process learns, but the fast process will forget how it adapted to the counter rotation, while the slow process still exhibits part of its adaptation from the long initial training, resulting in a rebound.

This model postulates the reaching behavior exhibited on trial t (X_t1_), is the sum of the output of the slow (X_s,t1_) and fast process (X_f,t1_) on the same trial:

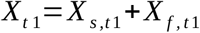

Both processes learn from errors on the previous trial (e_t0_) by means of a learning rate (L_s_ and L_f_), and they each retain some of their previous state (X_s,t0_ and X_f,t0_) by means of their retention rates (R_s_ and R_f_):

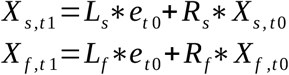

The model is further constrained by making sure the learning rate of the slow process is lower than that of the fast process: L_s_ < L_f_, and by having the retention rate of the slow process be larger than that of the fast process: R_s_ > R_f_. We constrained the parameters to the range [0,1].

All model fitting was done on the mean angular reach deviation at peak velocity during all training reaches, regardless of target angle. The error term was set to zero during the final error clamp phase of the experiment, as the participant did not experience any performance error. The model was fit in R 3.6.1 ^43^ using a least mean-squared error criterion on the six best fits resulting from a grid-search. The parameter values corresponding to the lowest MSE between data and model was picked as the best fit, and this was repeated for all groups.

### Rate of Change

We used an exponential decay function with an asymptote to estimate the rate of change for each of the three trial types. The value of each process on the next trial (p_t1_) is the current process’ value (p_t0_) minus the product of the rate of change (L) multiplied by the error on the current trial, which is the difference between the asymptote (A) and the process’ value on the current trial (p_t0_).

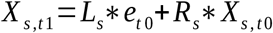

The parameter L was constrained to the range [0,1], and the parameter A to [0,2·max(data)]. For all groups, this model was fit to 1) the slow process from the two-rate model, and 2) the reach data. It was also fit to each group’s test trial data; 3) no-cursor reach deviations and both 4) active and 5) passive localizations. For the latter three kinds of fits, a zero was prepended to account for the fact that responses in these trials already changed through the previous training trial. The parameters were also bootstrapped (1k resamples per fit) across participants to get a 95% confidence interval for both parameters. The first trial where the modelled process based on the group average fell inside the bootstrapped confidence interval for the asymptote is taken as the saturation trial.

The datasets for the current study are available on Open Science Framework, https://osf.io/9db8v/, while the code and analysis scripts are available on github, https://github.com/JennR1990/TwoRateProprioception.

## Acknowledgements

We thank our two reviewers for their feedback, as it significantly improved the manuscript. This work was supported by a Canadian Network for Research and Innovation in Machining Technology NSERC Operating grant (DYPH) and the German Research Foundation (DFG) under grant no. HA 6861/2-1 (BMtH). The funders had no role in study design, data collection and analysis, decision to publish, or preparation of the manuscript.

